# Enhanced co-expression of cyclin F and USP7 in luminal A breast cancer correlates with endocrine resistance, high oxidative phosphorylation, and low inflammatory signatures

**DOI:** 10.64898/2026.01.08.698348

**Authors:** Savitha S. Sharma, P. S. Hari, R. Nivedhitha

## Abstract

Cyclin F is a non-canonical cyclin that functions as the substrate-recognition subunit of the SCF^Cyclin F^ E3-ubiquitin ligase complex. By targeting specific proteins for degradation, cyclin F plays an important role in proteostasis and genomic stability. We recently showed that cyclin F interacts with USP7, a deubiquitylating enzyme. USP7-mediated stabilization of ERα and PHF8 is linked to breast carcinogenesis. On the other hand, recent studies have implicated SCF^Cyclin F^ in modulating the CDK4/6-RB axis, a pathway that plays a crucial role in tumorigenesis and progression in the HR^+^/HER2^−^ breast cancer.

Herein, we performed an in-silico analysis to investigate the role of USP7-cyclin F axis in HR+/HER2-breast cancer. Cyclin F and USP7 transcripts were positively correlated across cancer cell lines and the luminal A breast cancer in the TCGA cohort. In both the METABRIC and TCGA breast cancer cohorts, high cyclin F-USP7 co-expression was synergistically associated with poor survival only in the endocrine-treated luminal A group, but not in those without endocrine treatment, suggesting an association with endocrine resistance. Notably, the transcriptomic profiles of high cyclin F-USP7 tumors were associated with signatures of endocrine resistance. Cancer Dependency Map analysis of CRISPR knockout of cyclin F or USP7 on cellular fitness suggested cyclin F influences the CDK4-RB pathway via RBL2 and/or PLK4, while USP7 might influence the ESR1-CDK4 network via HUWE1. Further, pathway analysis of differentially abundant proteins in the TCGA cohort identified enrichment of upregulated proteins in the oxidative phosphorylation and cell cycle pathways, and downregulated proteins in the immune pathways.

## Introduction

Breast cancer is the most common cancer in women, contributing to a significant number of deaths globally. It is broadly stratified into 4 major molecular subtypes – namely luminal A (encompassing hormone receptor-positive [HR+] and HER2-negative tumors), luminal B (HR+ that can be HER2^−^ or HER2^+^), HER2-enriched, and basal-like (often overlapping with the triple-negative breast cancer) [1]. Among these, the HR^+^/HER2^−^ subtype is the most prevalent, accounting for approximately 60-75% of reported cases in Western women and 50-60% in Indian women [2, 3]. Estrogen receptor-alpha (ERα) activation is a key driver of not only mammary gland development, but also of tumorigenesis and progression in the HR^+^/HER2^−^ breast cancer [4]. ERα knockout mice exhibit resistance to oncogene-induced malignant transformation underscoring the importance of ERα in breast tumorigenesis. Cyclin D1 is a known ERα-target gene. Furthermore, cyclin D1- or CDK4-deficient mice are refractory to various oncogene-triggered breast tumorigenesis [5, 6]. Due to this reliance on the estrogen-modulated ERα-Cyc D1/CDK4 signalling axis, patients with HR+ disease are typically treated with ER antagonists, such as tamoxifen, fulvestrant, or with aromatase inhibitors that suppress estrogen production, and in the metastatic setting with first line endocrine therapy combined with CDK4/6 inhibitors [7]. While most HR^+^/HER2^−^ breast cancers initially respond to endocrine therapy, 5-10% exhibit intrinsic resistance and 20-30% acquire resistance over a span of several years [8]. Multiple mechanisms of endocrine resistance have been uncovered, including, loss of ER expression, ER mutations enabling their ligand-independent activation, aberrant interactions between ER and its coactivators/corepressors, and genomic alterations in the PI3K/mTOR pathways driving hyperactivation of the cyclin D/CDK4/6-RB axis [8].

Cyclin F (encoded by CCNF), a cell-cycle regulated non-canonical cyclin, is the founding member of the F-box family of proteins. It functions as the substrate-recognition subunit of the SCF^Cyclin F^ (Skp1-Cul1-F box) E3 ubiquitin ligase complex, recruiting substrates to the E3, and enabling their ubiquitylation and subsequent degradation [9]. By targeting specific proteins for degradation, cyclin F plays a key role in proteostasis and genomic stability [10]. Intriguingly, recent studies have begun to implicate SCF^Cyclin F^ in modulating the CDK4/6-RB axis and G1/S transition [11]. Firstly, in response to mitogens, phosphorylated AKT stabilizes cyclin F and enhances SCF^Cyclin F^ complex formation [12]. Secondly, SCF^Cyclin F^ ubiquitylates and degrades p130 (RBL2), a pocket-protein closely related to RB that functions as a repressor of E2F-responsive genes [11]. By promoting the degradation of p130, cyclin F may enhance E2F-driven transcription and cell-cycle progression, potentially contributing to endocrine resistance mechanisms in HR+ breast cancer.

Adding another layer of regulation to the ERα-CDK4 axis is USP7, a deubiquitylating enzyme that stabilizes and/or alters the function of numerous oncogenic proteins by reversing their ubiquitylation [13]. Like cyclin F, USP7 is a master regulator of genomic stability. It is frequently overexpressed in multiple cancer, including breast cancer, and associated with tumor aggressiveness and therapy resistance [14]. USP7 was shown to interact with and stabilize ERα [15]. We recently showed that USP7 interacts with cyclin F and stabilizes it by protecting from proteasomal degradation [16]. We also identified USP7 as a positive regulator of cyclin F mRNA [16]. This finding positions USP7 as a critical upstream regulator of SCF^Cyclin F^. Dysregulation of the USP7-cyclin F axis could potentiate aberrant CDK4/6-RB signalling, potentially contributing to endocrine and/or CDK4/6 inhibitor resistance.

In this study, we performed an in-silico analysis to investigate the role of USP7-cyclin F regulatory axis in HR+/HER2-breast cancer. We show a significant positive correlation between cyclin F and USP7 mRNA expression across cancer cell lines and the luminal A breast cancer in the TCGA cohort. Furthermore, in both the METABRIC and TCGA breast cancer cohorts, high cyclin F-USP7 co-expression was synergistically associated with worse clinical outcomes only in the endocrine-treated luminal A group, but not in those without endocrine treatment, suggesting an association with endocrine resistance. Cancer Dependency Map (DepMap) analysis of loss-of-function of cyclin F or USP7 on cellular fitness suggested cyclin F might influence the CDK4-RB pathway by physically and/or genetically interacting with RBL2 and PLK4, while USP7 might influence the ESR1-CDK network via HUWE1. Further, pathway analysis of differentially abundant proteins in the TCGA cohort identified enrichment of upregulated proteins in the oxidative phosphorylation and cell cycle pathways, and downregulated proteins in the immune pathways. In addition, the transcriptomic profiles of high cyclin F-USP7 coexpressing tumors in both the TCGA and METABRIC cohorts were associated with signatures of endocrine resistance.

## Material and Methods

Analysis of DepMap datasets: Using the DepMap Data Explorer 2.0 tool, gene expression data were retrieved for cyclin F, USP7, and USP47 as log2-transformed TPM+1 from the Cancer Cell Line Encyclopedia (depmap.org, DepMap24Q4 release). Pearson correlation analysis was performed between the genes across the 1673 cancer cell lines (at the time of analysis). Correlation outputs (r and p value) and scatter plots were directly exported from the portal.

Similarly, to identify genes co-dependent with ESR1, CDK4, CCNF, USP7, or HUWE1, we interrogated the Achilles dataset (DepMap24Q4 release), which contains the gene-essentiality scores from the genome-wide CRISPR knockout screens across 1183 cell lines. The Pearson’s correlation coefficient of essentiality scores across 1183 cell lines and 17700 genes (at the time of analysis) was then analysed. Using Metascape (metascape.org), pathway enrichment analysis was performed for 0.5 % of the most highly correlated co-dependent genes for each of the aforementioned genes, across the following gene-sets and processes: GO Biological Processes, GO Molecular Function, Hallmark gene sets, Reactome gene sets, KEGG pathway, Wikipathways, and Canonical pathways. Venn diagrams were prepared using Venny 2.1.

### Analysis of TCGA datasets for CCNF and USP7 amplifications

Amplification frequency of CCNF and USP7 across various cancer types was retrieved from cBioPortal using the TCGA pan-cancer whole genome analysis study. An oncoprint was generated using cBioPortal to examine co-amplification of CCNF with USP7 in the Breast Invasive Carcinoma (TCGA, Firehose Legacy) cohort.

### Gene expression and Kalpan-Meier-survival analysis from METABRIC and TCGA datasets

The normalized TCGA-BRCA gene-expression data was downloaded from the UCSC Xena Cancer Genome Browser in RSEM normalised counts, which was then log_2_ +1 transformed. For METABRIC, normalized microarray expression data, clinical data was downloaded from cBioPortal. Disease-specific survival (DSS) information for the samples was obtained from Pereira *et al* [17]. The clinical and survival information for the TCGA samples were also obtained from the UCSC Xena database.

Normalised expression values for USP7 and CCNF was then extracted and samples from each cohort were divided into high and low expression groups by the median expression of each gene. A composite score was the prepared by averaging the normalized expression values of USP7 and CCNF. The median of this composite score was used to dichotomize patients into high-score and low-score groups for Kaplan–Meier survival analysis.

### Proteomics analysis from TCGA breast cancer datasets

Proteomics data for the samples used in the TCGA RNAseq analysis was obtained from Mertins et al [18]. Normalised log2 transformed proteomics abundances for both the CCNF-USP7 high and low groups were then used to compute the differentially abundant proteins

### Pathway and process enrichment analysis

The Metascape tool (metascape.org) was employed for pathway and process enrichment analysis as well as protein-protein interaction (PPI) enrichment analysis [19]. Pathway analysis was performed separately for the significant differentially upregulated and downregulated genes/proteins, with the following ontology sources: KEGG Pathway, GO Biological Processes, GO Molecular Functions, Reactome Gene Sets, Hallmark Gene Sets, Canonical Pathways, CORUM, WikiPathways, and PANTHER Pathway. Metascape calculates and clusters terms based on hypergeometric p-values and enrichment factor > 1.5 (the enrichment factor is the ratio between the observed counts and the counts expected by chance). The tool then performs hierarchical clustering on the enriched terms, and sub-trees with a similarity of > 0.3 were considered a cluster. The most statistically significant term within a cluster was chosen to represent the cluster.

For each list of differentially expressed genes or proteins, PPI enrichment analysis was carried out with the following databases: STRING, BioGrid, OmniPath, and InWeb_IM. Only physical interactions in STRING (physical score > 0.132) and BioGrid are used. The resultant network contained the subset of proteins that form physical interactions with at least one other member in the list. If the network contained between 3 and 500 proteins, the Molecular Complex Detection (MCODE) algorithm was applied to identify densely connected network components.

### Gene-set enrichment analysis

The WebGestalt tool (webgestalt.org) was applied to identify signatures of endocrine resistance in the transcriptomic profiles of CCNF-USP7 low and high groups. The gene symbols along with their log2FC values were uploaded to WebGestalt and analysed against a combined functional database comprised of human MSigDB hallmark (gsea-msigdb.org), and Creighton_endocrine_therapy_resistance 1 and 2 gene sets [20]. The normalised enrichment scores, along with p and FDR values were exported and used to determine enrichment of the signatures.

## Results

### Cyclin F and USP7 transcript levels positively correlate with each other

Our previous study showed that USP7 positively regulated cyclin F mRNA levels and that cyclin F also physically interacts with USP7. Genes that function in the same pathway are often coregulated or co-expressed. We, therefore, investigated the relationship between cyclin F and USP7 mRNA expression across 2 large datasets: 1673 cancer cell lines and the TCGA breast cancer cohort. Expression data were retrieved from the DepMap24Q4 release and expression values for cyclin F and USP7 were extracted as log2-transformed TPM+1. Pearson correlation analysis across the 1673 cancer cell lines revealed a significant positive correlation between cyclin F and USP7 (Pearson r = 0.633, p = 1.23 × 10^−187^; Figure 1A). On the other hand, the expression of cyclin F was weakly correlated with that of USP47, the most closely related DUB to USP7 (Pearson r = 0.269, p = 3.37 × 10^−29^; Figure 1B). These data suggest that cyclin F and USP7 may functionally interact with each other. Cyclin F and USP7 are co-amplified in 5-15% of breast invasive carcinoma (Suppl. figure 1A, B). We therefore examined whether the expression of the two genes was correlated among the breast cancer subtypes in the TCGA dataset (Figure 1 C-F). Among the 4 subtypes, cyclin F transcript levels showed a modest, significant positive correlation with that of USP7 in the luminal A and HER2-enriched subtypes, with a Pearson correlation of 0.54 (p < 2.2 × 10^−16^) and 0.5 (p < 1.9 × 10^−5^) respectively (Figure 1C, E). Overall, these findings suggest that the two proteins are co-regulated or participate in common pathways.

**Figure 1.**
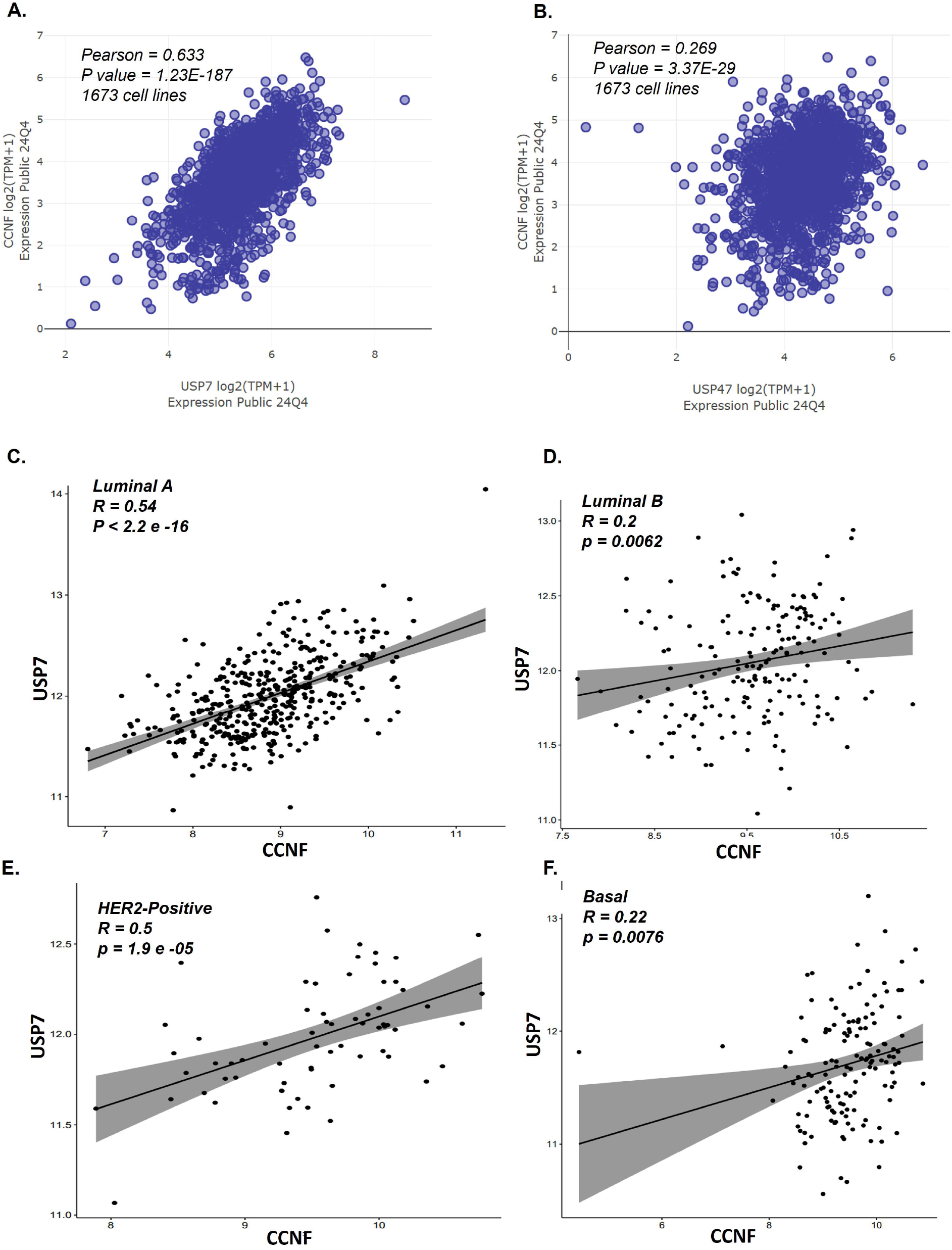
Cyclin F and USP7 transcript levels positively correlate with each other. **A and B.** Pearson’s correlation analysis between CCNF and USP7, or CCNF and USP47 gene expression levels across the 1673 cancer cell lines from the Cancer Cell Line Encyclopedia using DepMap Data Explorer 2.0. **C-F:** Normalised transcript levels for CCNF and USP7 were plotted for each of the breast cancer subtypes from the TCGA dataset. The Pearson’s correlation coefficient (r) between expression levels for the two genes, along with their p values, are indicated for each subtype.

### High co-expression of cyclin F-USP7 correlates with shortened survival and endocrine resistance

To assess the prognostic significance of high co-expression of USP7 and cyclin F in the luminal A breast cancer subtype, we performed Kaplan-Meier survival analysis using the METABRIC dataset. Patients were stratified based on co-expression of both the genes, using a median cutoff to identify cyclin F-USP7-high- and low-expressors, and disease-specific survival (DSS) was analysed. In the subset of patients with luminal A subtype who received endocrine therapy, high co-expression of cyclin F and USP7 was significantly associated with worse DSS (p = 0.00069; Figure 2C). While high cyclin F or USP7 expression, individually, also associated with DSS (p values of 0.07 and 0.02, respectively), the association with DSS was more significant in tumors with high combined cyclin F-USP7 expression (Figure 2A, B). More importantly, in luminal A patients not treated with endocrine therapy, no significant association was observed between cyclin F-USP7 co-expression and survival outcomes (p = 0.31; Figure 2D). This suggests that the prognostic impact of this co-expression pattern is treatment specific and may reflect endocrine resistance mechanisms. Furthermore, the observed prognostic impact was specific for cyclin F-USP7 co-expression, because replacement of cyclin F with cyclin A2, a cyclin that is most closely related to cyclin F, did not exhibit a significant association with DSS (Figure 2E, F).

**Figure 2.**
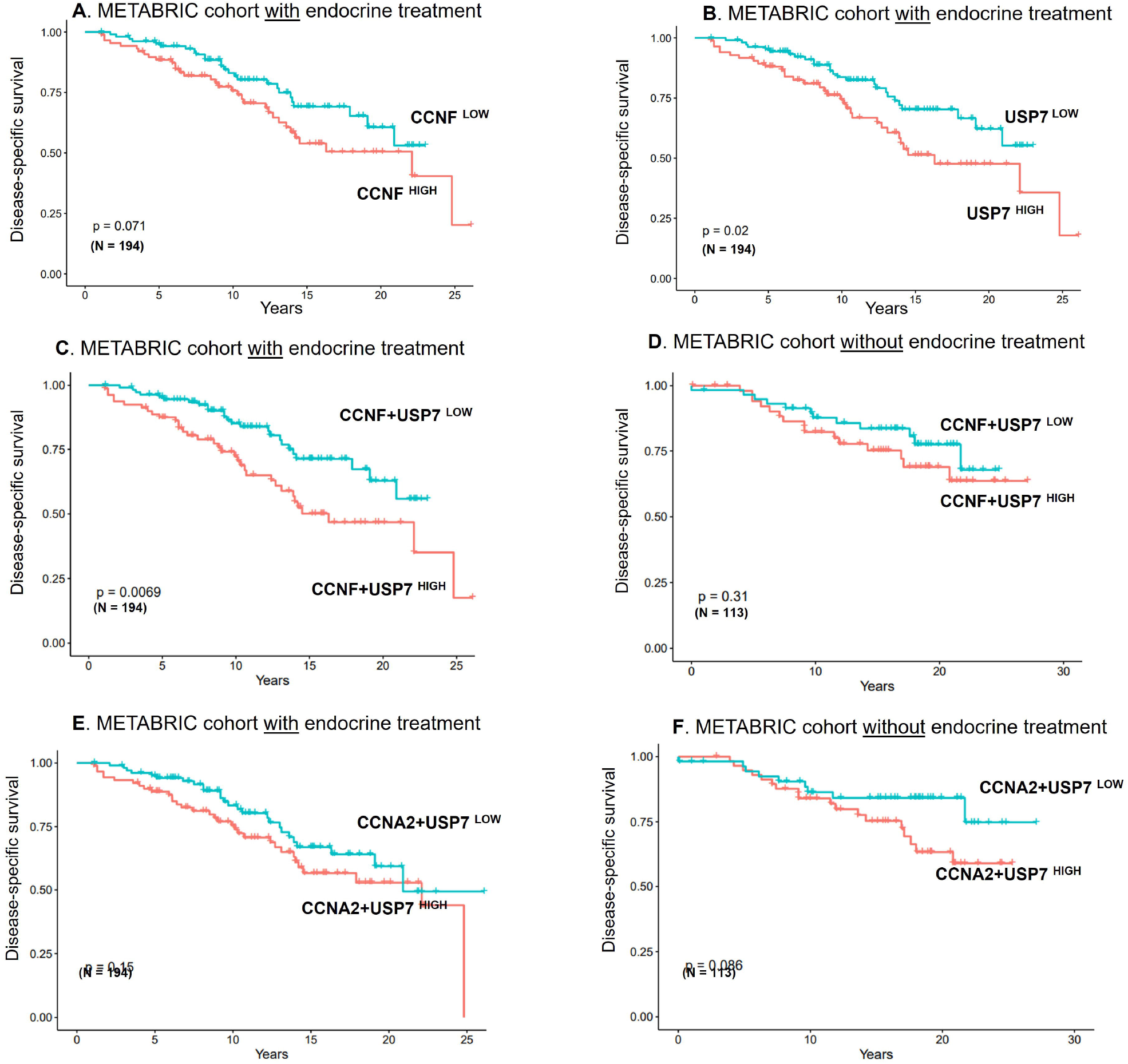
Correlation of gene expression levels with disease-specific survival (DSS) in the METABRIC cohort. **A.** Kaplan-Meier (KM) analysis of patients with luminal A breast cancer, who received endocrine therapy, stratified based on CCNF^High^ or CCNF^Low^ expression. **B**. KM analysis of patients with luminal A breast cancer, who received endocrine therapy, stratified based on USP7^High^ or USP7^Low^ expression. **C**. KM analysis of patients with luminal A breast cancer, who received endocrine therapy, stratified based on CCNF+USP7^High^ or CCNF+USP7^Low^ co-expression. **D**. KM analysis of patients with luminal A breast cancer, without endocrine treatment, stratified based on CCNF+USP7^High^ or CCNF+USP7^Low^ co-expression. **E**. KM analysis of patients with luminal A breast cancer, who received endocrine therapy, stratified based on CCNA2+USP7^High^ or CCNA2+USP7^Low^ co-expression. F. KM analysis of patients with luminal A breast cancer, without endocrine treatment, stratified based on CCNA2+USP7^High^ or CCNA2+USP7^Low^ co-expression. The p values and sample size (N) are indicated for each plot.

Similar to the METABRIC cohort, Kaplan-Meier analysis using the TCGA breast cancer dataset also showed that high co-expression of cyclin F and USP7 mRNA was significantly associated with worse progression-free survival (PFS; p = 0.048) in patients with luminal A subtype who received endocrine therapy (Figure 3C). Again, in this cohort too, no significant association was observed between PFS and expression of cyclin F (p = 0.16) or USP7 (p = 0.14) individually (Figure 3A, B).

**Figure 3.**
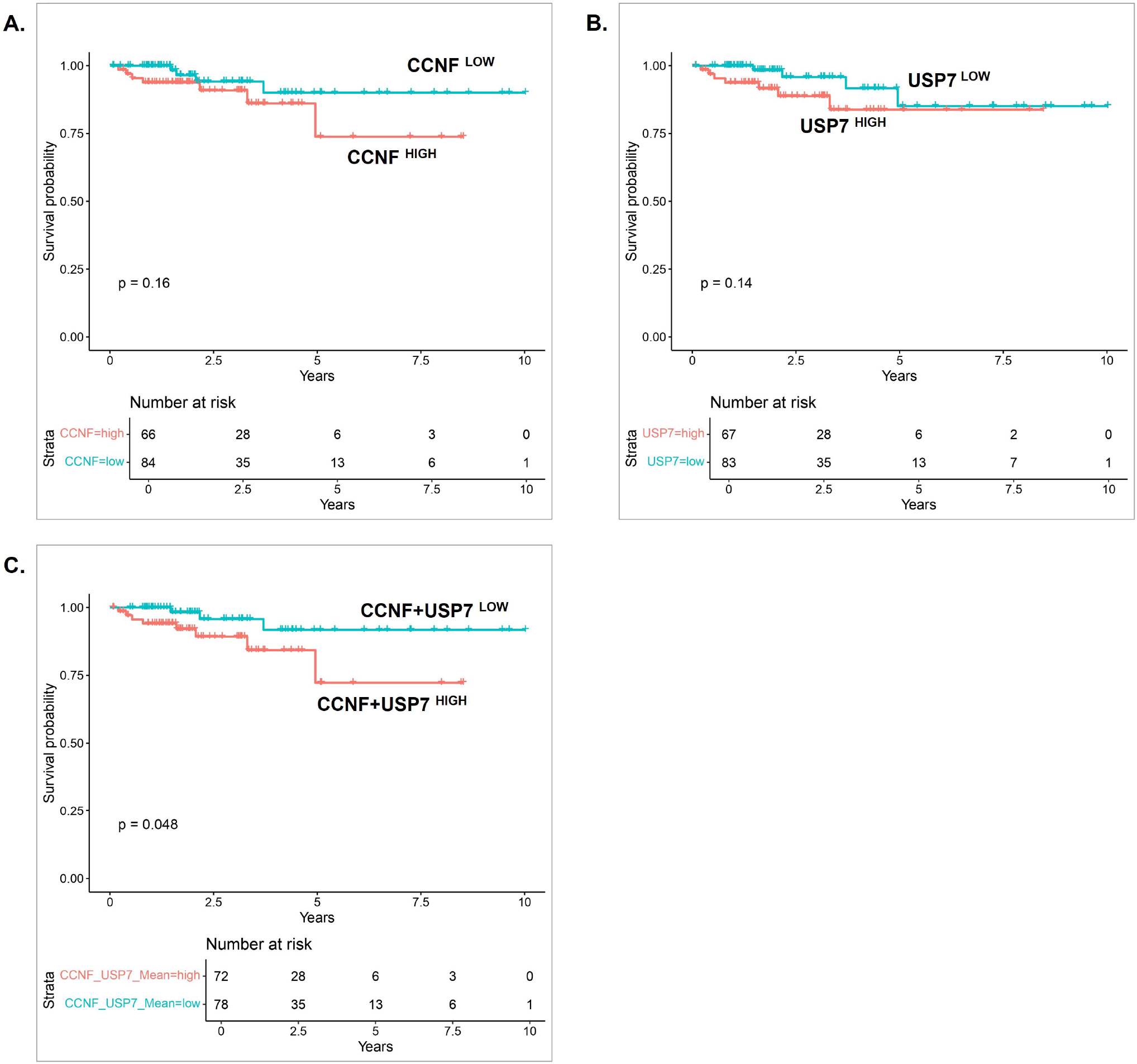
Correlation of gene expression levels with progression-free survival (PFS) in the TCGA cohort. **A.** Kaplan-Meier (KM) analysis of patients with luminal A breast cancer, who received endocrine therapy, stratified based on CCNF^High^ or CCNF^Low^ expression. **B**. KM analysis of patients with luminal A breast cancer, who received endocrine therapy, stratified based on USP7^High^ or USP7^Low^ expression. **C**. KM analysis of patients with luminal A breast cancer, who received endocrine therapy, stratified based on CCNF+USP7^High^ or CCNF+USP7^Low^ co-expression. The p values and sample size (N) are indicated for each plot.

### Cyclin F and USP7 fitness scores highly correlate with that of genes in the CDK4-RB and ESR1-CDK4 networks

Using the DepMap dataset, Emanuele and colleagues had previously reported that the cyclin F fitness correlates with that of several members of the CDK4-RB network, namely CDK4, CDK6, CCND1/ Cyclin D1, RBL1, and RBL2 [11]. Further experiments demonstrated that cyclin F physically interacts with RBL2/p130, targeting it for degradation and promoting G1-S phase transition [11]. To gain insights into the relationship between cyclin F, USP7, and endocrine resistance, and uncover how the two genes might physically and/or genetically interact with the ESR1-CDK4 axis, we utilized the updated DepMap dataset. We first analysed the Pearson’s correlation coefficient for CDK4 knockout across 1183 cell lines and 17700 genes (at the time of analysis). Among the top 1% of the genes whose fitness scores were most highly correlated with CDK4 knockout were CCNF (27^th^ position) and USP7 (116^th^ position), along with CCND1, FOXA1, SPDEF, HUWE1, and ESR1 (Figure 4A; Suppl. file 1). Likewise, among the top 0.5% of the highly correlating genes for ESR1 knockout were FOXA1, SPDEF, CDK4, CCND1, and HUWE1 (Figure 4C; Suppl. file 1). These results show that ESR1 and cyclin D1-CDK4 pathways not only functionally interact with each other, but also that cyclin F and USP7 might influence these pathways.

**Figure 4.**
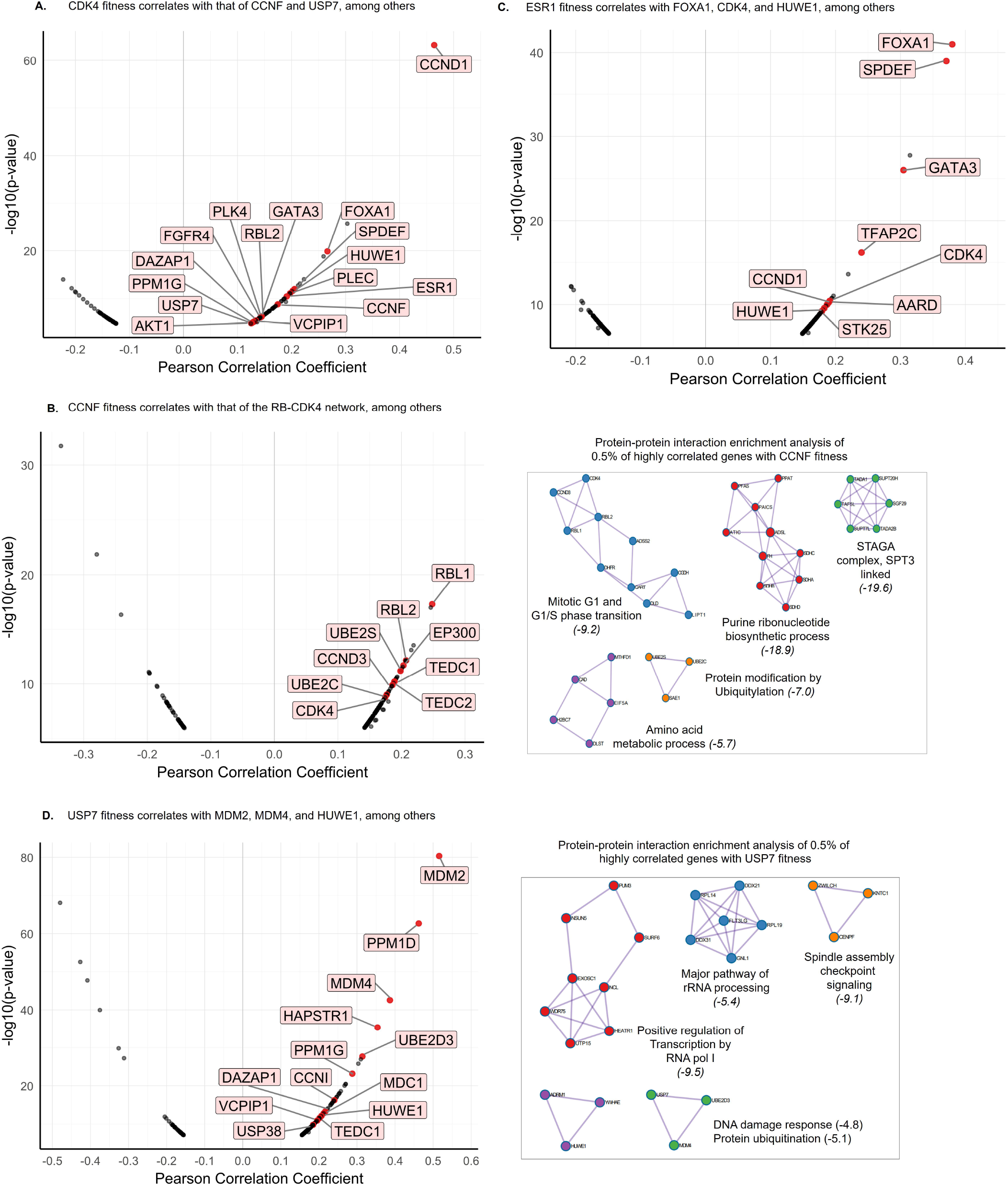
DepMap data analysis of Co-dependent genes for CDK4, ESR1, CCNF, and USP7: **A.** Volcano plot depicting the top 1% of the genes whose fitness scores were significantly correlated with that of CDK4. **B**. Volcano plot depicting the top 1 % of the genes whose fitness scores were significantly correlated with that of CCNF, along with PPI enrichment analysis of the positively correlated genes (panel right of B; Log10(P) are indicated in brackets). **C**. Volcano plot depicting the top 1 % of the genes whose fitness scores were significantly correlated with that of ESR1. **D**. Volcano plot depicting the top 1 % of the genes whose fitness scores were significantly correlated with that of USP7, along with PPI enrichment analysis of the positively correlated genes (panel right of D; Log10(P) are indicated in brackets).

To understand how cyclin F and USP7 might synergize with the ESR1-CDK4 axis, we performed similar analysis of the DepMap dataset. Pearson’s correlation coefficient analysis for CCNF knockout identified RBL1, RBL2, CCND3, and CDK4 among the top 0.5 % of highly correlated genes (Figure 4B; Suppl. file 1). PPI enrichment analysis of the top 0.5% of these highly correlated genes identified pathways associated with mitotic G1 and G1/S phase transition, purine nucleotide biosynthetic process, amino acid metabolic process, protein modification by ubiquitylation, and STAGA complex SPT3 linked pathways (Figure 4B).

For USP7 knockout, MDM2/4, PPM1D/G, CCNI, and HUWE1 were among the top 0.5% of the highly correlated genes (Figure 4D; Suppl. file 1). PPI enrichment analysis of the top 0.5% of the codependent genes showed strong enrichment in pathways including that of ribosome biogenesis, rRNA processing, mitotic spindle checkpoint signalling, and DNA-damage checkpoint, and protein ubiquitylation (Figure 4D). Venn diagram analysis indicated that CDK4 was common between ESR1- and CCNF-codependent genes (Figure 5A); likewise, HUWE1 was common between ESR1− and USP7-codependent genes (Figure 5A). Additional analysis indicated that CCNF might functionally interact with CDK4 via RBL2 and/or PLK4 (Figure 5B). On the other hand, USP7 is likely to functionally influence CDK4 via HUWE1, MDM2, PPM1G, DAZAP1, C4ORF46, and USP38 (Figure 5B; Suppl. file 1). Among these USP7-codependent genes, HUWE1 knockout fitness most correlated with ESR1, FOXA1, SPDEF, CDK4, and USP7 networks, among others (Figures 5C, D; Suppl. file 1). Collectively, these data suggest that cyclin F might influence the ESR1-CDK4 axis via RBL2 and/or PLK4, while USP7 might influence it via HUWE1.

**Figure 5.**
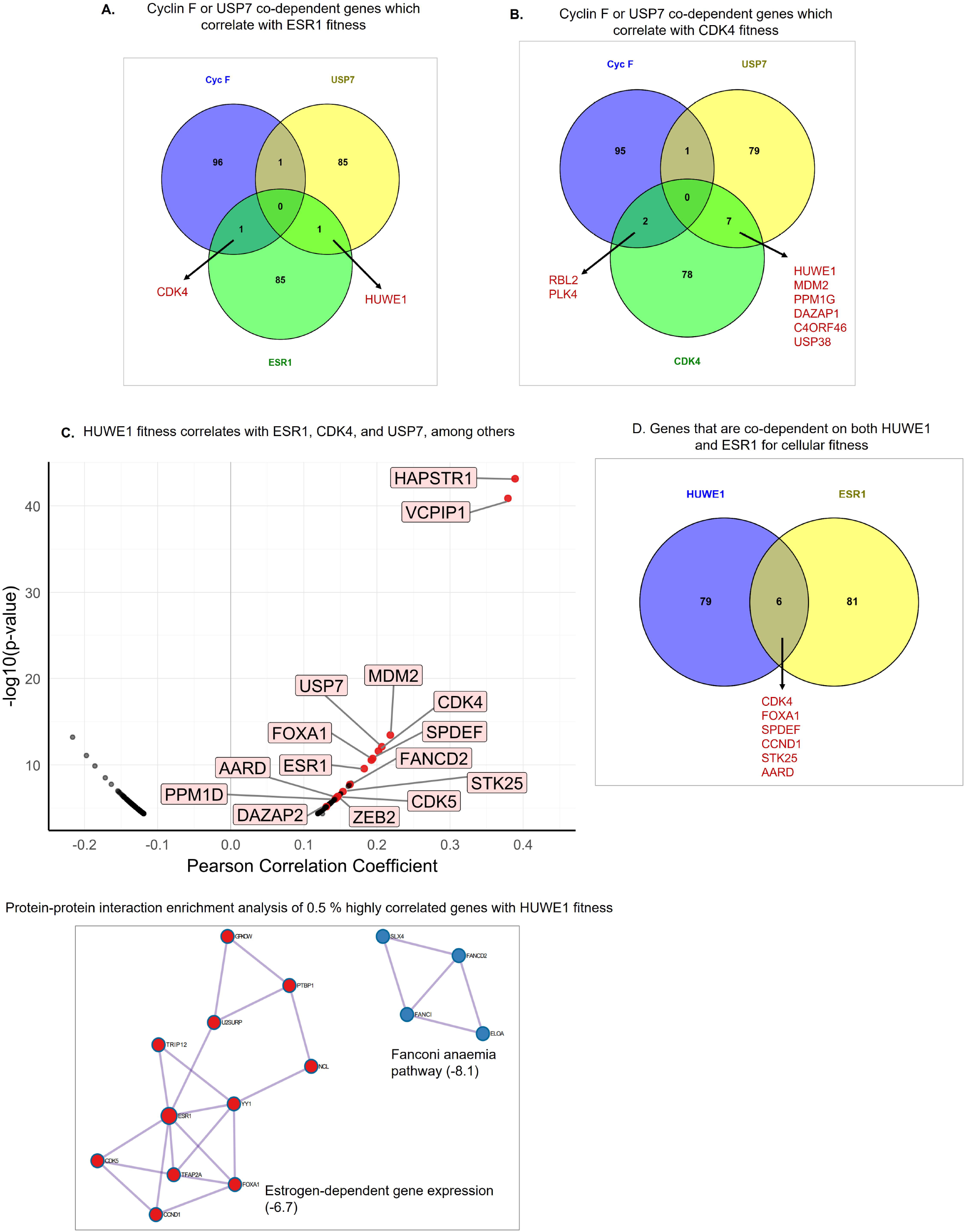
CCNF or USP7 co-dependent genes that are required for ESR1 or CDK4 fitness. **A.** Venn diagrams showing CCNF or USP7 co-dependent genes that correlate with ESR1 fitness in the DepMap Achilles dataset. **B**. Venn diagrams showing CCNF or USP7 co-dependent genes that correlate with CDK4 fitness. **C**. Volcano plot depicting the top 1 % of the genes whose fitness scores were significantly correlated with that of HUWE1. PPI enrichment analysis of the positively correlated genes from the HUWE1 volcano plot (panel below C; Log10(P) are indicated in brackets). D. Venn diagram depicting genes that are co-dependent on both HUWE1 and ESR1 for cellular fitness.

### Differentially abundant proteins in high cyclin F-USP7 co-expressing tumors

Since cyclin F and USP7are known to play a key role in regulating protein stability, we anticipated a more immediate and direct impact of cyclin F-USP7 co-expression on differential protein expression. To examine the differentially abundant proteins between cyclin F-USP7 high- and low-expressors, we analysed the proteomics dataset from the TCGA breast cancer cohort, available through the CPTAC initiative. We compared tumors with high co-expression of cyclin F-USP7 versus those with low co-expression in the endocrine-treated luminal A subtype and identified 444 upregulated genes (log2FC > 0.5 and p ≤ 0.05), 184 downregulated (log2FC < −0.5 and p ≤ 0.05) (Suppl. file 2). Of note, USP7 protein was significantly upregulated in the high cyclin F-USP7 group; cyclin F protein, on the other hand, was undetectable in any of the samples in the dataset (Suppl. file 2). Pathway enrichment analysis of the upregulated proteins revealed significantly enriched for pathways related to oxidative phosphorylation, mitochondrial gene expression, mitochondrial organisation, mitochondrial protein degradation, metabolism of RNA, E2F targets, mitotic cell cycle, base excision repair, DNA replication and retinoblastoma gene in cancer, among others (Figure 6A). On the other hand, the downregulated proteins were significantly enriched for pathways related to neutrophil degranulation, metal sequestration by antimicrobial proteins, positive regulation of apoptotic process, complement, coagulation, and regulation of cell projection organization (Figure 6B).

**Figure 6.**
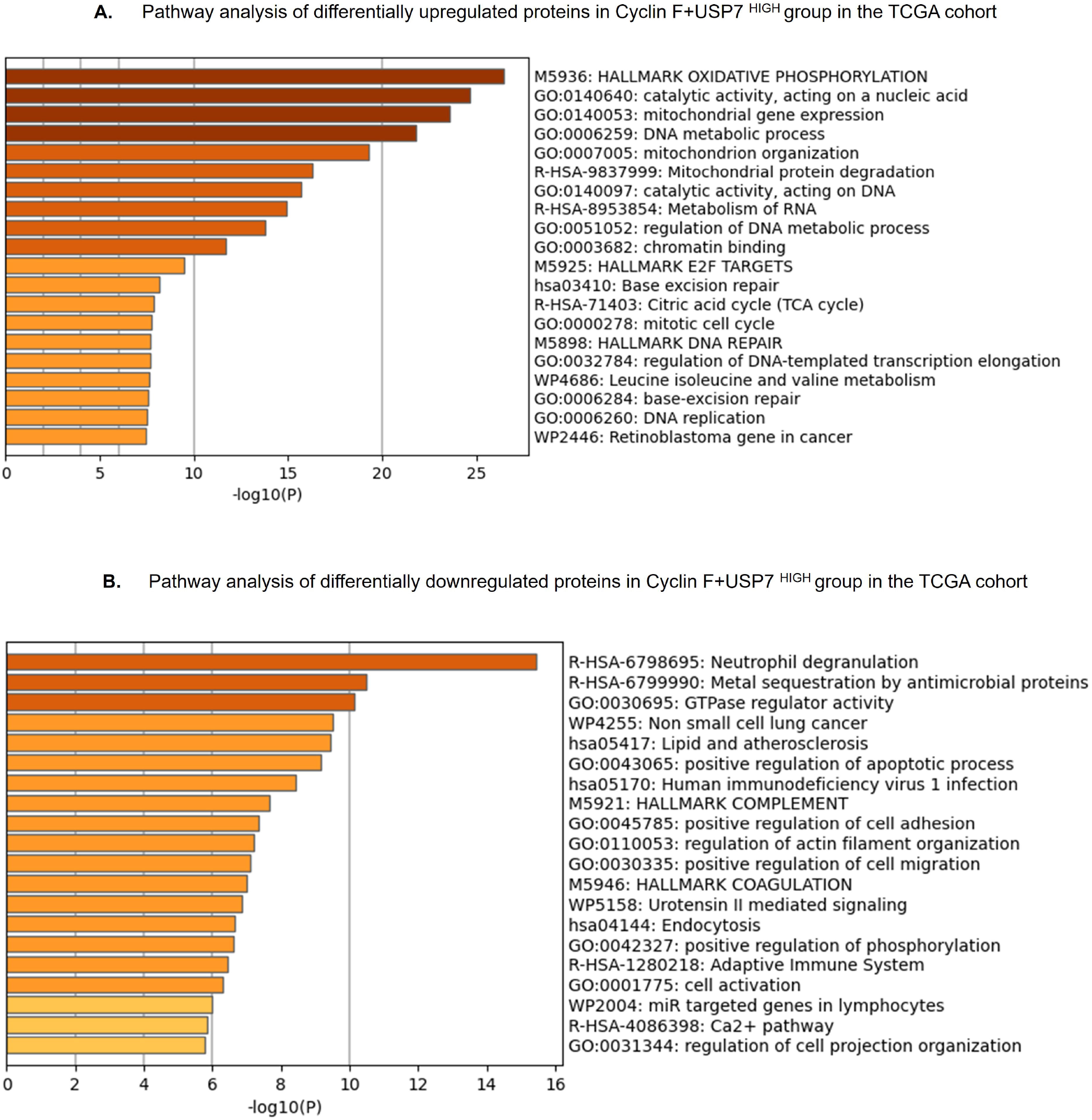
Pathway analysis of differentially abundant proteins in the TCGA cohort. **A.** Pathway analysis of the differentially upregulated proteins in CCNF+USP7^HIGH^ tumors from patients with luminal A breast cancer, who received endocrine therapy, using METASCAPE. **B**. Pathway analysis of the differentially downregulated proteins in CCNF+USP7^HIGH^ tumors.

### Differentially expressed genes in high cyclin F-USP7 co-expressing tumors

To investigate the transcriptional program associated with high co-expression of cyclin F and USP7, we performed a differential gene expression analysis, comparing tumors with high co-expression of these genes versus those with low co-expression in the endocrine-treated luminal A subtype of both the METABRIC and TCGA cohorts. In the METABRIC cohort, we identified 74 upregulated genes (log2FC > 0.4 and p ≤ 0.05), 234 downregulated (log2FC < −0.4 and p ≤ 0.05) (Suppl. file 3). In the TCGA cohort, we identified 161 upregulated genes (log2FC > 0.5 and p ≤ 0.05) and 379 downregulated (log2FC < −0.5 and p ≤ 0.05) (Suppl. file 4). The upregulated genes, in the METABRIC cohort, were significantly enriched for pathways related to estrogen response-early, mammary gland alveolus development, female pregnancy, and hormone metabolic process (Figure 8A). Similarly, the upregulated genes, in the TCGA cohort, were significantly enriched for pathways related to mitotic cell cycle, epithelial cell differentiation, estrogen response-early, and glycolysis, among others (Figure 7A). The downregulated genes, in the METABRIC cohort, were significantly enriched for pathways related to TNFA signalling via NFκB, NABA core matrisome, epithelial mesenchymal transition, negative regulation of cell population proliferation, and immune effector process among others (Figure 8B). On the other hand, the downregulated genes in the TCGA cohort, were significantly enriched for pathways related to TNFA signalling via NFKB, inflammatory response, positive regulation of cell adhesion, and response to steroid hormone, among others (Figure 8B). Importantly, GSEA analysis of the transcriptional profiles of cyclin F-USP7-high tumors from both the cohorts were associated with signatures of endocrine resistance (Figure 7C, 8C). Collectively, these results indicate that high cyclin F-USP7 co-expression is associated with upregulation of the estrogen early-response, mitotic cell cycle, and epithelial differentiation pathways, and downregulation of NF-κB and inflammatory networks.

**Figure 7.**
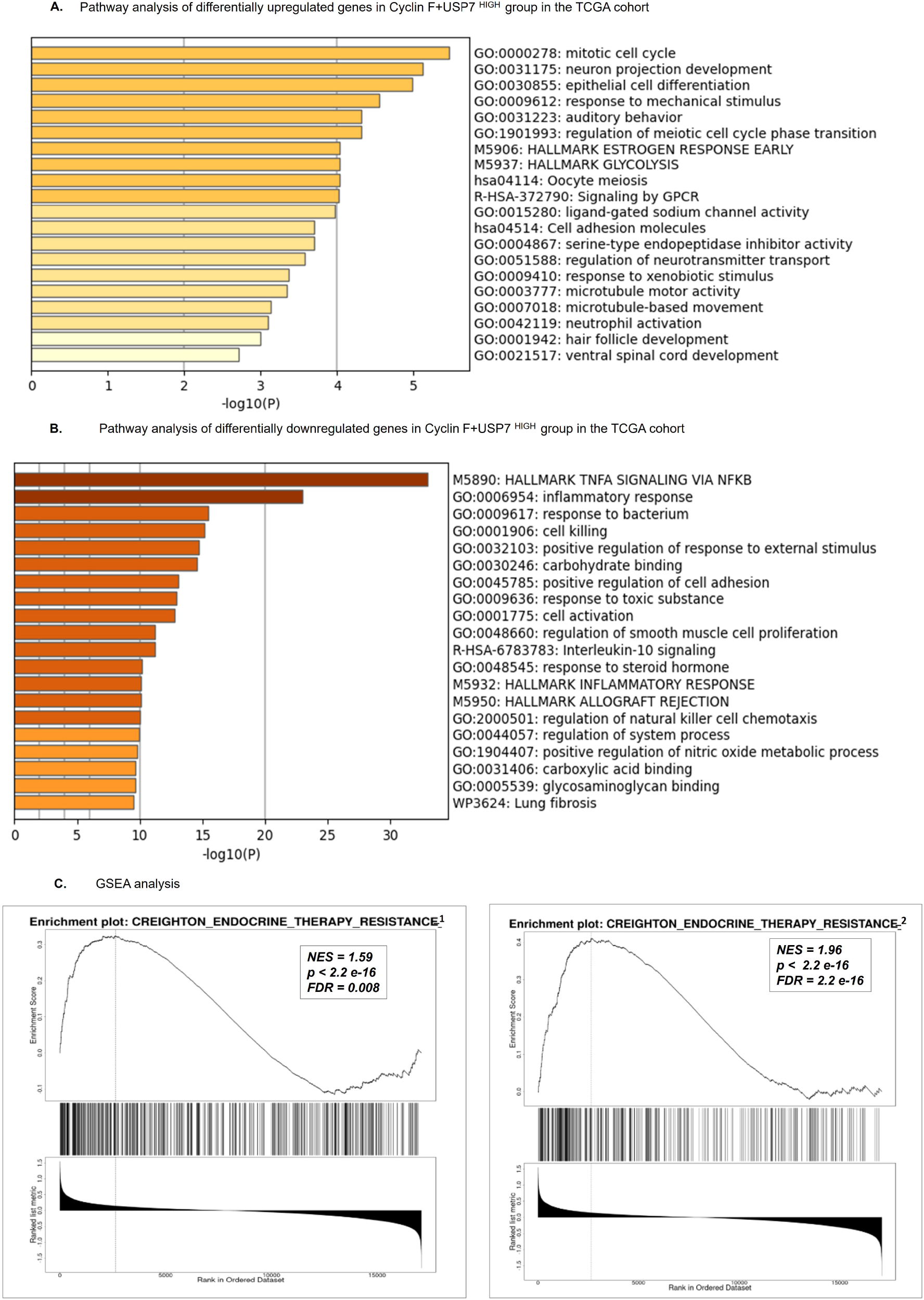
Pathway analysis of differentially expressed genes in the TCGA cohort. **A.** Pathway analysis of the differentially upregulated genes in CCNF+USP7^HIGH^ tumors from patients with luminal A breast cancer, who received endocrine therapy, using METASCAPE. **B**. Pathway analysis of the differentially downregulated genes in CCNF+USP7^HIGH^ tumors. C. GSEA analysis of the ranked gene expression list from CCNF+USP7^HIGH^ tumors examining signatures of endocrine resistance.

**Figure 8.**
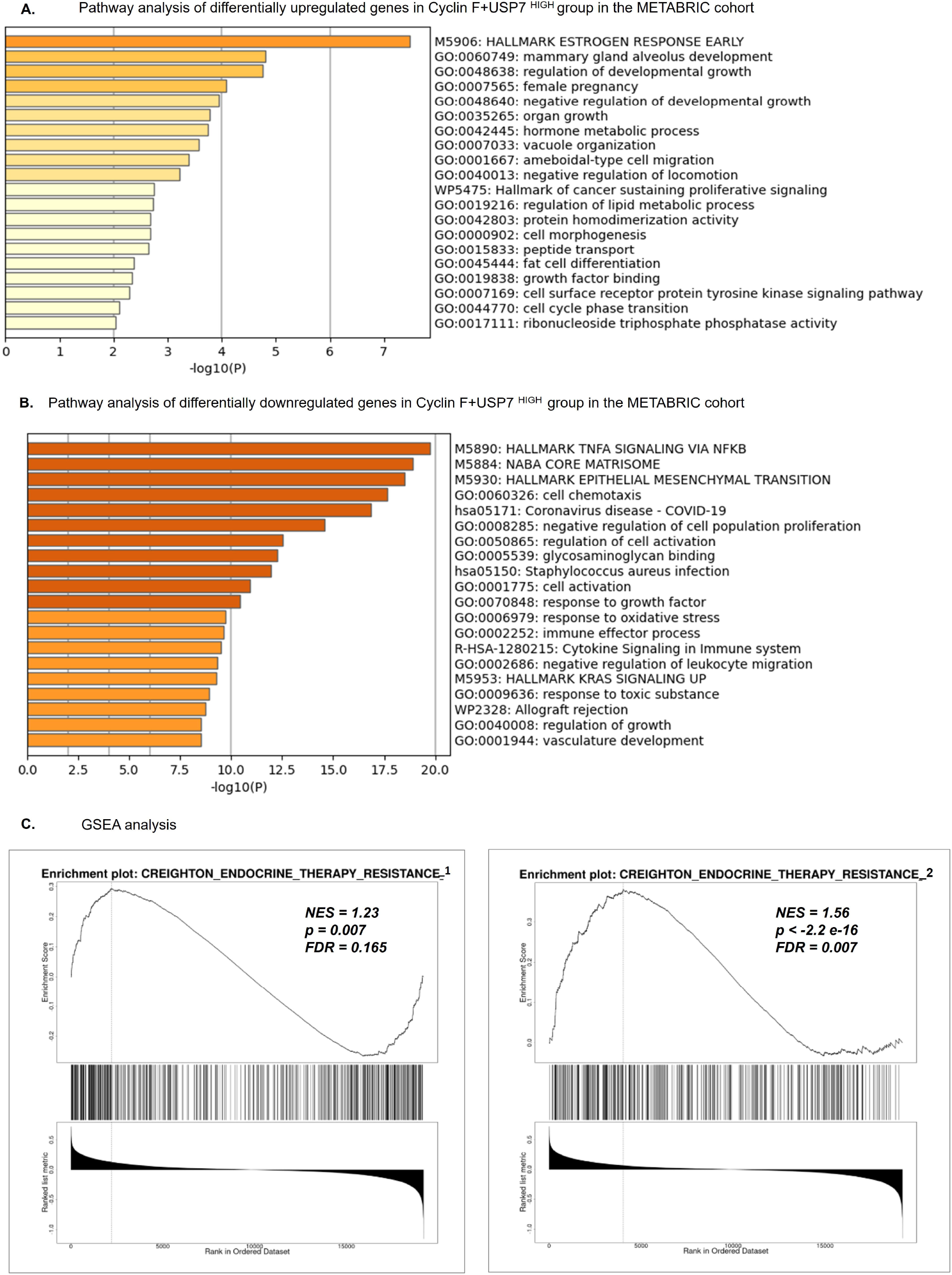
Pathway analysis of differentially expressed genes in the METABRIC cohort. **A.** Pathway analysis of the differentially upregulated genes in CCNF+USP7^HIGH^ tumors from patients with luminal A breast cancer, who received endocrine therapy, using METASCAPE. **B**. Pathway analysis of the differentially downregulated genes in CCNF+USP7^HIGH^ tumors. **C**. GSEA analysis of the ranked gene expression list from CCNF+USP7^HIGH^ tumors examining signatures of endocrine resistance.

## Discussion

Cyclin F is an E3-ubiquitin ligase, while USP7 is a deubiquitylase. Whereas cyclin F is a cell-cycle regulated protein, USP7 is not. Importantly, both proteins are key regulators of genomic stability, with well-documented roles in DNA replication, cell-cycle checkpoint, DNA repair, and transcriptional regulation [10, 13, 21]. Cyclin F expression, at both mRNA and protein levels, is upregulated in clear-cell renal cell carcinoma and breast cancer, and associates with poor survival [22, 23]. In gliomas and hepatocellular carcinoma, cyclin F is downregulated and appears to function as a tumor suppressor [24, 25]. USP7, likewise, has context dependent roles in cancer. For instance, its overexpression in cancers of the lung, breast, gastric, colon, cervix, and others are known to associate with drug resistance and metastasis [13]. Our in-silico analyses suggests that high cyclin F-USP7 co-expression correlates with shortened DSS or PFS only in the endocrine-treated group of luminal A subtype of breast cancer, but not in those without endocrine treatment. Further, such an association was either weak or absent when the cohort was stratified based on cyclin F or USP7 expression alone. Collectively, these observations suggest that the two proteins might synergistically contribute to endocrine resistance.

The DepMap CRISPR knockout datasets for cyclin F or USP7 provided potential mechanistic insights into how the two genes might synergistically contribute to endocrine resistance. Our analyses reveal that cyclin F is likely to influence the ESR1-CDK4 axis by physically/genetically interacting with RBL2 and/or PLK4. Indeed, Emanuele and colleagues showed that cyclin F can ubiquitylate and target RBL2/p130 for degradation, resulting in activation of the DREAM complex and E2F-target genes [11]. How cyclin F might influence PLK4 is not known. Of note, upregulated PLK4 in the context of p53 inactivation (probably mediated, in the study cohort, by upregulated USP7 and HUWE1) can contribute to endocrine resistance [26]. On the other hand, analysis of the USP7 co-dependent genes revealed that USP7 might influence ESR1-CDK4 axis via interaction with HUWE1, in addition to influencing CDK4 via PPM1G, DAZAP1, C4ORF46, and USP38. For example, USP7 itself is known to downregulate p53 protein levels by stabilizing MDM2, thereby reducing p21 and removing the brake on CDK4/6 activity [27]. Notably, loss of p53 is associated with resistance to CDK4/6 inhibitor in HR+/HER2− breast cancers [28]. Additionally, USP7 might influence the ESR1-CDK4 axis via HUWE1, a HECT-domain-harboring E3 ligase implicated in the regulation of diverse pathways, including DNA-damage response, proliferation, autophagy, and transcriptional control [29]. Importantly, DepMap analysis identified co-dependency of HUWE1 with both ESR1 and CDK4 pathways; in particular, PPI network analysis of the HUWE1-codependent genes identified MCODE functional networks related to estrogen-dependent gene expression and TFAP2 (AP-2) family regulates transcription of growth factors and their receptors (Figure 5C). HUWE1 is often found overexpressed in cancers of breast, lung, prostate, colon, stomach, multiple myeloma, and uterus, where it associates with disease progression and might function as an oncogene [30]. In brain tumors, HUWE1 is underexpressed, and high expression is associated with better survival outcome [29]. Adhikary *et al* demonstrated that HUWE1-mediated ubiquitylation of c-Myc is required for transactivation of its target genes and is essential for tumor cell proliferation [31]. Importantly, c-Myc is known to transactivate CDK4, on one hand, and repress many INK and CIP1/KIP1 inhibitor expression, on the other, leading to enhanced cyclin D/CDK4 activity [29]. These findings suggest that upregulation of HUWE1 might lead to endocrine resistance.

As members of the ubiquitination pathway, cyclin F and USP7 play crucial roles in regulating protein stability, localization, and/or function. Both proteins are known to physically interact with each other [16]. The interplay between the opposing E3-ligase activity of cyclin F and the deubiquitylase activity of USP7 is expected to have an immediate effect at the level of protein abundance and activity of key cyclin F and/or USP7 substrates, including transcription factors, chromatin modifiers, and cell-cycle regulators. Such protein changes will likely have consequent cascading effects on gene transcription. Therefore, to gain insights into the relevant biological pathways associated with endocrine resistance in high cyclin F-USP7 tumors, we first examined the differentially abundant proteins using the TCGA proteomics dataset. First, high CCNF-USP7 mRNA expression correlated with upregulated levels of USP7 protein compared to controls. Cyclin F protein, on the other hand, was undetectable in any of the samples in the dataset, probably because cyclin F is a short-lived protein, hence low abundant, and below detection limit in mass-spectrometry based proteomics. Thus, while we do not know the cyclin F protein levels in our TCGA cohort, using immunohistochemistry, Lui *et al* have previously reported higher levels of cyclin F protein in all subtypes of breast cancer compared to normal tissue [23]. Furthermore, among the differentially upregulated proteins in cyclin F-USP7-high tumors, we found significant enrichment of proteins related to oxidative phosphorylation, E2F targets, and DNA repair pathway. Conversely, the differentially downregulated proteins were statistically enriched for pathways related to neutrophil degranulation, metal sequestration by antimicrobial proteins, apoptosis, coagulation, and adaptive immune system, among others. The upregulation of E2F target genes could in part be fuelled by cyclin F-mediated RBL2 degradation, as reported by others and our DepMap analysis [11]. Alternatively, higher cyclin F level in these tumor cells is able to effectively counteract the pro-apoptotic activities caused by E2F hyperactivation [32, 33]. Recently, El-Botty *et al* reported high oxidative phosphorylation in bone-metastasis-derived and CDK4/6i-resistant PDX models of ER+ breast cancer [34]. Notably, our DepMap analysis identified many genes of the TCA cycle pathway, including, SDHA, SDHB, SDHC, SDHD, and FH as CCNF co-dependent genes, suggesting that cyclin F might play a role in regulation of oxidative phosphorylation (Suppl. file 1). With respect to USP7, among its numerous substrates/interactors, we found HUWE1, FOXA1, YY1, PPM1G, UHRF2, DDR1, and ERCC6 to be upregulated at the protein levels in the cyclin F-USP7-high tumors [13]. Of these, HUWE1 (discussed earlier), along with FOXA1 and YY1/2 might contribute to the reprogramming of the ERα transcriptional output and endocrine resistance [35, 36]. Indeed, GSEA analysis of the transcriptional profiles of cyclin F-USP7-high tumors were associated with signatures of endocrine resistance (Figure 7C, 8C). Notably, these tumors were characterized by downregulation of TNFA signalling via NFκB, inflammatory response and complement pathways, suggesting low-inflammatory phenotypes (Figures 7B, 8B).

In conclusion, our in-silico analysis points to a synergistic role for cyclin F and USP7 in endocrine resistance. We propose that cyclin F likely contributes to endocrine resistance by modulating the G1-S progression, while USP7 does so by influencing the ESR1 transcriptional program via HUWE1. A major limitation of this study is that it is purely correlational and does not determine causal relationships between high cyclin F-USP7 co-expression and endocrine resistance. Thus, further validation of these findings is warranted in independent cohorts using immunohistochemistry and transcriptional analyses, along with other cell-based experiments to assess functional interaction of cyclin F and USP7 with the ESR1-CDK4 axis.

## Supporting information

Supplementary File 1_DepMap analyses

Supplementary File 2_TCGA_DAPs

Supplementary File 3_METABRIC_DEGs

Supplementary File 4_TCGA_DEGs

## Acknowledgements

We thank Guruguhan Sivakumar for preparation of the volcano plots and Paturu Kondaiah for critical review of the manuscript. This work is supported in part by the Lady Tata Memorial Trust – Institutional Research Grant (Mumbai, India) to S.S.S.

## Author contributions

S.S.S designed the study, performed DepMap, METASCAPE, and GSEA analyses, interpreted all results, wrote the manuscript. H.P.S. performed transcriptomic, proteomic, and survival analyses using TCGA and METABRIC breast cancer datasets, interpreted the results, and reviewed the manuscript. N.R wrote the manuscript with S.S.S, and prepared the figures. All authors read and approved of the manuscript.

## Figure Legends

Supplemental Figure 1: **A**. Bar graph showing amplification frequency of CCNF and USP7 across various cancer types (TCGA pan-cancer cohort) as retrieved from cBioPortal. **B**. Oncoprint showing frequency of the indicated gene alterations in the Breast Invasive Carcinoma (TCGA, Firehose Legacy) cohort, as retrieved from cBioPortal.

