## Supplementary figures and images for "Enhanced co-expression of cyclin F and USP7 in luminal A breast cancer correlates with endocrine resistance, high oxidative phosphorylation, and low inflammatory signatures"

### Supplementary Figure 1

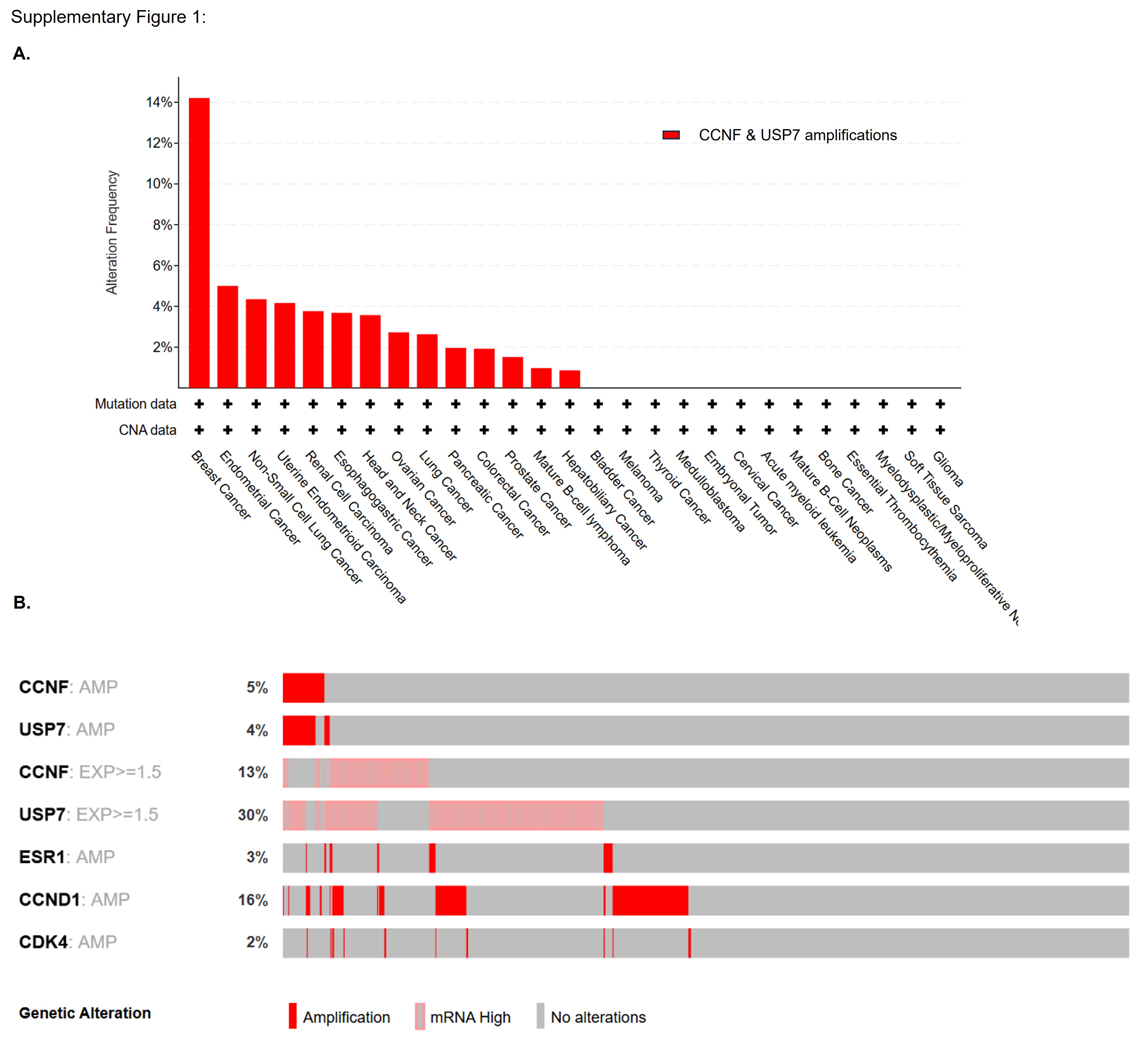
